# Identification of “missing links” in C- and D-ring cleavage of steroids by *Comamonas testosteroni* TA441

**DOI:** 10.1101/2023.02.02.526765

**Authors:** Masae Horinouchi, Toshiaki Hayashi

## Abstract

*Comamonas testosteroni* TA441 degrades steroids aerobically via aromatization and cleavage of the A- and B-ring, followed by D- and C-ring cleavage via b-oxidation. We previously characterized most of the above degradation steps; however, a few intermediate compounds remained unaccounted for. We hypothesized that cleavage of the D-ring at C13-17 required the ScdY hydratase and was followed by cleavage of the C-ring via the ScdL1L2 transferase. The reaction was expected to produce 6-methyl-3,7-dioxo-decane-1,10-dioic acid-Coenzyme A (CoA) ester. To verify this hypothesis, we constructed a plasmid that enabled targeted gene induction in TA441 mutant strains. The major product of ScdL1L2 was found to be 3-hydroxy-6-methyl-7-oxo-decane-1,10-dioic acid-CoA ester; whereas the substrate of ScdY was revealed to be a geminal diol, 17-dihydroxy-9-oxo-1,2,3,4,5,6,10,19-octanorandrost-8(14)-en-7-oic acid-CoA ester. This finding suggests that ScdY catalyzes the addition of a water molecule at C14 of 17-dihydroxy-9-oxo-1,2,3,4,5,6,10,19-octanorandrost-8(14)-en-7-oic acid-CoA ester, leading to D-ring cleavage at C13-17. The C9 ketone of the D-ring cleavage product is then converted to a hydroxyl group, followed by C-ring cleavage to produce 3-hydroxy-6-methyl-7-oxo-decane-1,10-dioic acid-CoA ester. Precise bacterial bile acid degradation pathway will be one of the key to investigate the gut–microbiota–brain axis, which affects human health and disease.

**IMPORTANCE:** Studies on bacterial steroid degradation were initiated more than 50 years ago primarily to obtain substrates for steroid drugs. The role of steroid-degrading bacteria in relation to human health is attracting growing attention. *Comamonas testosteroni* TA441 is the leading bacterial model of aerobic steroid degradation and the overall pathway has been outlined previously. However, a few intermediate compounds in C- and D-ring cleavage processes have remained unknown. Here, we identified the missing compounds and can now propose the complete A-, B-, C-, and D-ring cleavage pathway employed by steroid-degrading bacteria. This finding will facilitate the application of such microorganisms for the synthesis of specific steroid derivatives.

## INTRODUCTION

Steroid compounds have various function in both plants and animals, including humans where they participate in hormone, cholesterol, and cholic acid production. Studies on bacterial steroid degradation were initiated more than 50 years ago primarily to obtain materials for steroid drugs. The Actinobacterium *Rhodococcus equi* (formerly *Nocardia restrictus*) and Proteobacterium *Comamonas testosteroni* (formerly *Pseudomonas testosteroni*) are known for their ability to degrade steroid compounds. The underlying mechanism was extensively studied around 1960, which led to identification of the major intermediates in A- and B-ring degradation processes, and revealed a similar steroid degradation pathway in both bacteria (1–5). Steroids are attracting increasing attention because of their influence on human health in relation to pathogenic bacteria. In *Mycobacterium tuberculosis* H37Rv, the *mce4* operon encoding a cholesterol import system is essential for bacterial persistence in the lungs of chronically infected animals and for growth within interferon-gamma-activated macrophages (6). Cholesterol catabolism and its broader utilization by *M. tuberculosis* are important for pathogen maintenance in the host (7). Cholic acid and deoxycholic acid are synthesized from cholesterol in the liver, from where they are secreted into the bile and then in the intestine. There, these primary bile acids are converted to secondary bile acids and other derivatives by intestinal bacteria, thereby affecting human health.

Genetic studies on steroid degradation by *C. testosteroni* started around 1990, and the enzymes catalyzing the early steps of this process, 17β-dehydrogenase (8–11), 3α-dehydrogenase (12–16), 3-oxo-Δ5-steroid isomerase (17, 18), Δ1-dehydrogenase (19), Δ4-dehydrogenase (20), and a positive regulator (21) were identified. However, the enzymes involved in steroidal ring cleavage remained unclear. The presently known mechanism employed by TA441 to degrade the four basic steroidal rings (A-, B-, C-, and D-rings) is summarized in Fig. 1. TA441 is the leading model of bacterial aerobic steroid degradation. It first converts steroids (e.g., testosterone, cholic acid, and their derivatives) to androsta-1,4-diene-3,17-dione (R1, R2 = H) or the corresponding derivative 7α,9α-dihydroxy-androsta-1,4-diene-3,17-dione (in cholic acid degradation) (compound **I**). Subsequent aromatization of the A-ring, ring cleavage, and hydrolysis lead to 2-hydroxyhexa-2,4-dienoic acid (**VI**) and 9,17-dioxo-1,2,3,4,10,19-hexanorandrostan-5-oic acid (**VII**) or the corresponding derivative 7α,9α-dihydroxy-17-oxo-1,2,3,4,10,19-hexanorandrostan-5-oic acid (in cholic acid degradation) (22–29). In the next set of reactions, which proceed by β-oxidation, coenzyme A (CoA) is first incorporated into **VII** by the CoA transferase ScdA (30), and the cleaved B-ring in **VII**-CoA ester is removed to produce 9α-hydroxy-17-oxo-1,2,3,4,5,6,10,19-octanorandrostan-7-oic acid **(XII**)-CoA ester (31–33). The C-ring is then dehydrogenated to 9,17-dioxo-1,2,3,4,5,6,10,19-octanorandrost-8(14)-en-7-oic acid (**XIV**)-CoA ester (34), followed by D-ring cleavage with the CoA hydratase ScdY. Based on the compounds isolated from a culture of TA441 with a mutated form of the C-ring cleavage enzyme ScdL1L2, 9-oxo-1,2,3,4,5,6,10,19-octanor-13,17-secoandrost-8(14)-ene-7,17-dioic acid (**XV**)-CoA ester, 13-hydroxy-9-oxo-1,2,3,4,5,6,10,19-octanor-13,17-secoandrost-8(14)-7,17-dioic acid (**XVI**)-CoA ester, and 9-hydroxy-1,2,3,4,5,6,10,19-octanor-13,17-secoandrost-13-ene-7,17-dioic acid (**XVII**)-CoA ester were proposed as products of D-ring cleavage (35). However, how these three compounds might be involved in the degradation process remained unclear. Our latest study suggested that 14-hydroxy-9-oxo-1,2,3,4,5,6,10,19-octanor-13,17-secoandrostane-7,17-dioic acid (**XVIII**)-CoA ester was the substrate of ScdL1L2 (36). The ring-cleavage product of **XVIII**-CoA ester is expected to be 6-methyl-3,7-dioxo-decane-1,10-dioic acid (**XIX**)-CoA ester, with molecular weight (MW) of 244. While we detected a small amount of **XVIII** in the ScdL1L2 mutant, we failed to identify any possible product of ScdL1L2 with MW of 244. Finally, one should not discard the possibility of unidentified intermediates arising from the process of D-ring cleavage. For example, addition of a water molecule to C14 of the **XIV**-CoA ester by an enoyl-CoA hydratase, would lead to the cleavage of the C8-C14 bond on the C-ring. In this study, we constructed a plasmid for inducible expression of target genes in TA441 and investigated the “missing links” in C- and D-ring cleavage.

**Fig 1.**
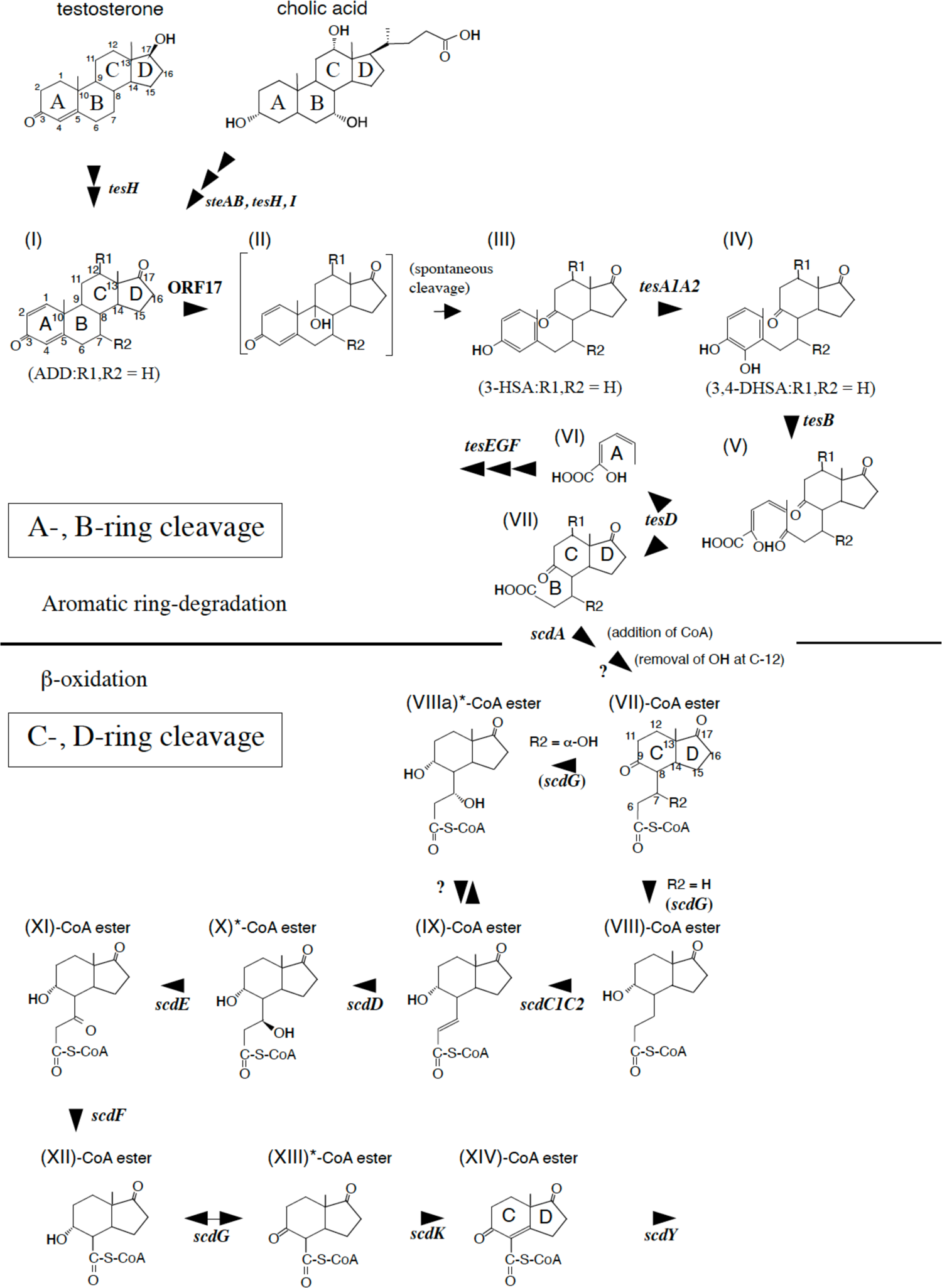
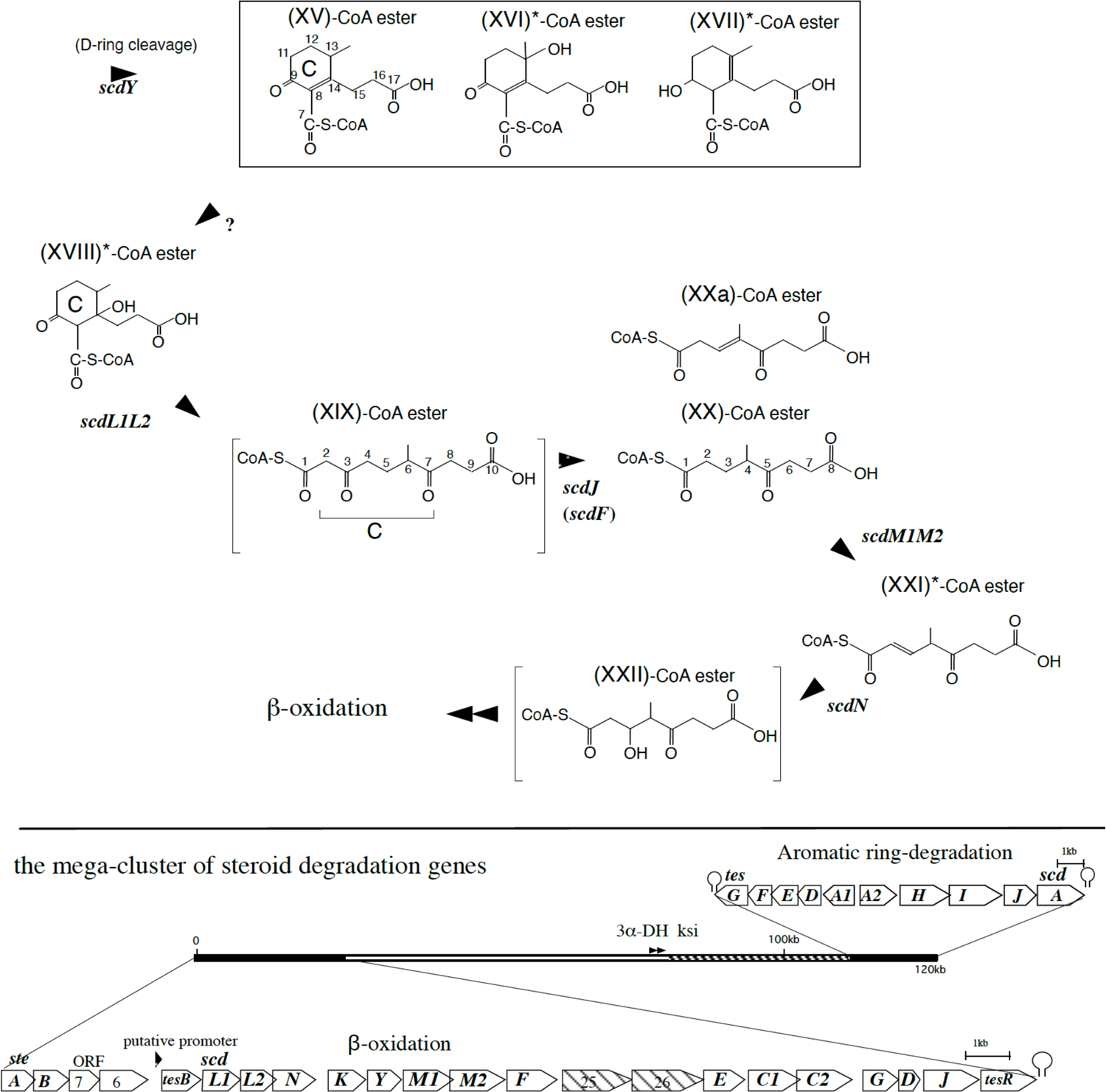
Steroid degradation pathway of *Comamonas testosteroni* TA441 revealed in our previous studies. Intermediate compounds were isolated and identified with NMR and high-resolution MS (HRMS) analysis (in β-oxidation process, compounds without CoA were identified) except for; compounds with * were identified with LC/MS/MS and experimentally confirmed, and compounds in square brackets are speculation. Compounds in the open box are possible intermediate compounds predicted from compounds identified in ScdL1L2-disrupted mutant culture. The mega-cluster of steroid degradation genes in *C. testosteroni* TA441 is shown below the degradation pathway; the aromatic ring-degradation gene cluster (*tesG* to *scdA*) and the β-oxidation gene cluster (*steA* to *tesR*) locate both ends of the 120kb-mega cluster. 3α-Hydroxy-dehydrogenase (3α-DH) gene and 3-ketosteroid Δ4-5 isomerase (ksi) gene are in the DNA region between the two clusters. Possible degradation genes for the side chain of cholic acid at C17 are found in the striped region. Compounds are; androsta-1,4-diene-3,17-dione (ADD) (R1,R2=H), (**I**); 9-hydroxy-androsta-1,4-diene-3,17-dione (ADD) (R1,R2=H), (**II**); 3-hydroxy-9,10-secoandrosta-1,3,5(10)-triene-9,17-dione (3-HSA) (R1,R2=H), (**III**); 3,4-dihydroxy-9,10-secoandrosta-1,3,5(10)-triene-9,17-dione (3,4-DHSA) (R1,R2=H), (**IV**); 4,5-9,10-diseco-3-hydroxy-5,9,17-trioxoandrosta-1(10),2-dien-4-oic acid (R1,R2=H), (**V**); (2*Z*,4*Z*)-2-hydroxyhexa-2,4-dienoic acid, (**VI**); 9,17-dioxo-1,2,3,4,10,19-hexanorandrostan-5-oic acid (3aα-*H*-4α [3′-propionic acid]-7aβ-methylhexahydro-1,5-indanedione) (R1,R2=H), (**VII**); 9α-hydroxy-17-oxo-1,2,3,4,10,19-hexanorandrost-6-en-5-oic acid, (**VIII**); 9α,7α-dihydroxy-17-oxo-1,2,3,4,10,19-hexanorandrost-6-en-5-oic acid, (**VIIIa**); 9-hydroxy-17-oxo-1,2,3,4,10,19-hexanorandrost-6-en-5-oic acid (**IX**); 9α,7β-dihydroxy-17-oxo-1,2,3,4,10,19-hexanorandrost-6-en-5-oic acid, (**X**); 9α-hydroxy-7,17-dioxo-1,2,3,4,10,19-hexanorandrost-6-en-5-oic acid, (**XI**); 9α-hydroxy-17-oxo-1,2,3,4,5,6,10,19-octanorandrostan-7-oic acid, (**XII**); 9,17-dioxo-1,2,3,4,5,6,10,19-octanorandrostan-7-oic acid, (**XIII**); 9,17-dioxo-1,2,3,4,5,6,10,19-octanorandrost-8(14)-en-7-oic acid, (**XIV**); 9-oxo-1,2,3,4,5,6,10,19-octanor-13,17-secoandrost-8(14)-ene-7,17-dioic acid, (**XV**); 13-hydroxy-9-oxo-1,2,3,4,5,6,10,19-octanor-13,17-secoandrost-8(14)-ene-7,17-dioic acid, (**XVI**); 9-hydroxy-1,2,3,4,5,6,10,19-octanor-13,17-secoandrost-13-ene-7,17-dioic acid, (**XVII**); 14-hydroxy-9-oxo-1,2,3,4,5,6,10,19-octanor-13,17-secoandrostane-7,17-dioic acid, (**XVIII**); 6-methyl-3,7-dioxo-decane-1,10-dioic acid, (**XIX**); 4-methyl-5-oxo-octane-1,8-dioic acid (**XX**); 4-methyl-5-oxo-oct-3-ene-1,8-dioic acid, (**XXa**); 4-methyl-5-oxo-oct-2-ene-1,8-dioic acid, (**XXI**); and 3-hydroxy-4-methyl-5-oxo-octane-1,8-dioic acid (**XXII**). Enzymes are; SteA (dehydrogenase for C12α-OH to C12-ketone), SteB (hydrogenase for C12-ketone to C12β-OH), TesH (Δ1-dehydrogenase), TesJ (**I-**hydroxylase at C9), TesA1A2 (**III-**hydroxylase at C4), TesB (*meta*-cleavage enzyme for **IV**), TesD (**V-**hydrolase), TesE (**VI-**hydratase), TesF (aldolase), TesG (acetoaldehyde dehydrogenase), ScdA (CoA-transferase for **VII**), ScdG (hydrogenase primarily for C9-OH of **XII-**CoA ester), ScdC1C2 (dehydrogenase for **VIII-**CoA ester at C6), ScdD (**IX-**CoA ester hydratase), ScdE (dehydrogenase for **X-**CoA ester at C7), ScdF (**XI-**CoA ester thiolase), ScdK (dehydrogenase for **XIII-**CoA ester at C8(14)), ScdY (likely to be **XIV-**CoA ester hydratase, but not confirmed yet), ScdL1L2 (CoA-transferase/isomerase involved in C-ring cleavage), and ScdN (**XXI-**CoA ester hydratase).

## RESULTS AND DISCUSSION

### Characterization of “missing link” intermediates in C- and D-ring cleavage

Our previous studies indicated that the enoyl-CoA hydratase ScdY and putative CoA transferase ScdL1L2 (Fig. 1) were responsible for cleaving the D- and C-ring of steroid compounds, respectively (34, 35). **XIV** (MW 208) was isolated from an ScdY-deficient mutant (ScdY^-^), suggesting that **XIV**-CoA ester was the substrate for ScdY. ScdL1L2 is homologous to the IpdAB hydrolase in *M. tuberculosis* H37Rv, which has been reported to cleave the C-ring of **XV**-CoA ester to produce **XIX**-CoA ester in the presence of a CoA transferase such as ScdF of TA441 (37) (Fig. 1). However, we were unable to detect **XIX** or its derivatives, even when using an ScdJ^-^ mutant, which lacked the CoA transferase required by the product of C-ring cleavage. A BLAST search has revealed that ScdL1L2 shows homology to the CoA transferase AtoDA (38) and SugarP_isomerase superfamily proteins. **XV** (MW 226), decarboxylated derivatives of **XVI** (MW 242), and **XVII** (MW 228) were identified by nuclear magnetic resonance and high-mass analysis in ScdL1^-^L2^-^ mutants (35). However, subsequent studies suggested that these compounds could arise during the isolation procedure, which was carried out under acidic conditions. Even though **XVIII**-CoA ester was suggested as the substrate of ScdL1L2 (36), the expected product, **XIX** (MW 244), was not detected by high-performance liquid chromatography with UV detection (samples were analyzed without extraction with ethyl acetate under acidic conditions) or ultraperformance liquid chromatography-mass spectrometry. Therefore, here, we cultured ScdY^-^ and ScdL1^-^L2^-^ cells with cholic acid and analyzed the cultures using reverse-phase liquid chromatography with tandem mass spectrometry (LC-MS/MS) (Fig. 2). In the ScdY^-^ culture, several overlapping peaks with *m/z* 207 were detected from retention time (RT) = 5.0 min to RT = 8.1 min (Fig. 2A). Diluting the sample 1/10 allowed to better distinguish these peaks (Fig. 2B). The one at RT = 8.1 min was identified as **XIV** based on comparison with known values (34); whereas the one at RT = 5.0 min was a fragment of a peak with *m/z* 225 (Fig. 2C). This finding implied that the peak with *m/z* 207 at RT = 5.0 min was derived from **XIV** following addition of a water molecule. The mass spectrum of the peak with *m/z* 225 at RT = 5.0 min revealed four major fragments (*m/z* 225, 207, 163, and 137); instead, that of the peak with *m/z* 207 at RT = 5.0 min revealed only three major fragments (*m/z* 207, 163, and 135), which corresponded to those of **XIV** (Fig. 3A–C). A water molecule can attack the C3 ketone, C17 ketone or C8(14) double bond on **XIV**. Here, the C17 geminal diol was converted automatically to **XIV** or **XV**, and the mass spectrum of **XV** identified the dominant fragment as *m/z* 137 (Fig. 3D). These results indicate that the compound with *m/z* 225 at RT = 5.0 min was 17-dihydroxy-9-oxo-1,2,3,4,5,6,10,19-octanorandrost-8(14)-en-7-oic acid (**XXIII**) (Fig. 3). The mass chromatograms of the ScdL1^-^L2^-^ culture revealed only compounds already identified in previous studies (Fig. 2D–G).

**Fig 2.**
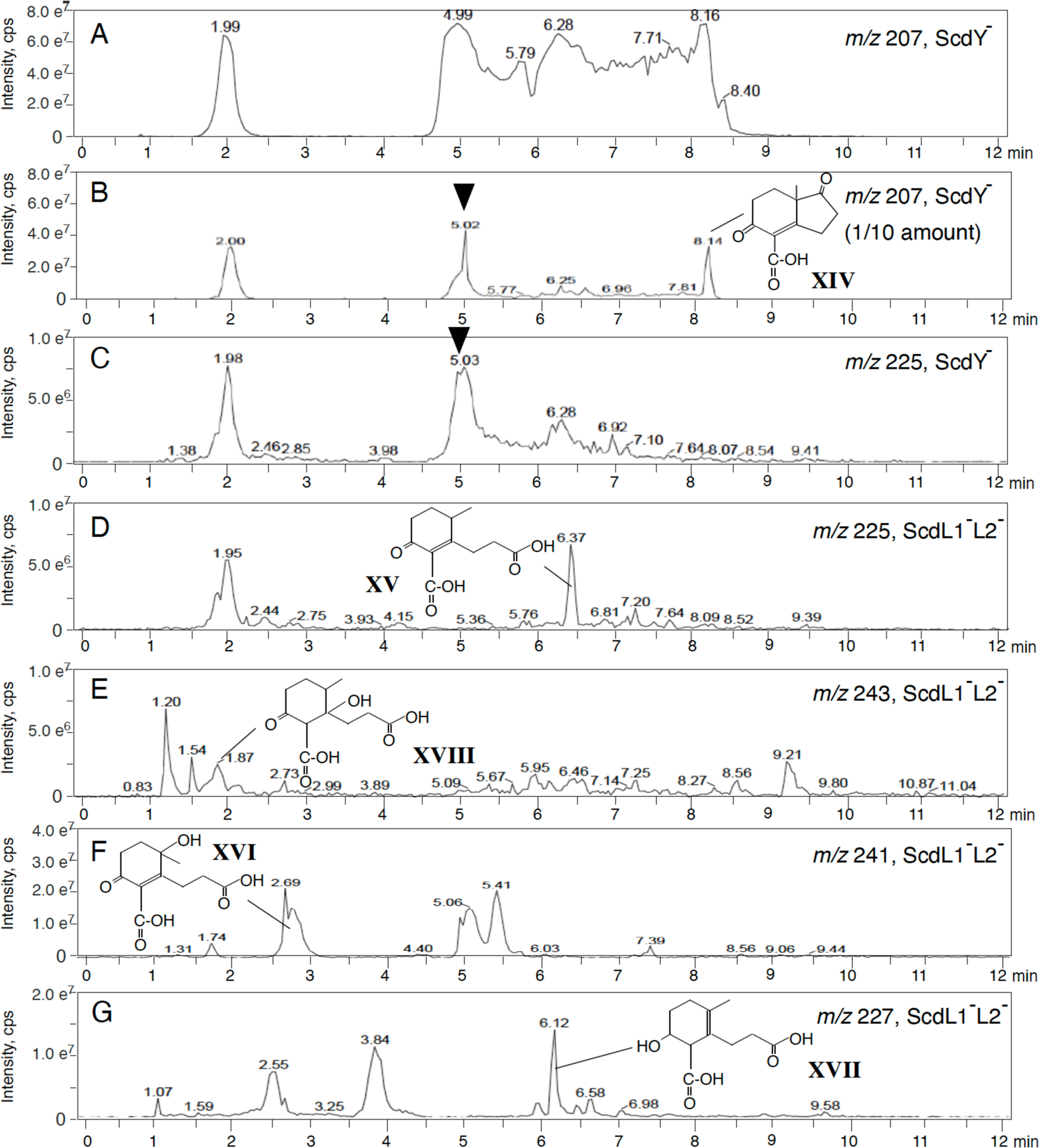
LC/MS/MS analysis of the culture of the ScdY^-^ and ScdL1^-^L2^-^ mutants. Panels indicate the mass chromatogram of *m/z* 207 (ScdY^-^) (A), 1/10 amount of (A) (B), *m/z* 225 (ScdY^-^) (C), *m/z* 225 (ScdL1^-^L2^-^) (D), *m/z* 243 (ScdL1^-^L2^-^) (E), *m/z* 241 (ScdL1^-^L2^-^) (F), and *m/z* 227 (ScdL1^-^L2^-^) (G). The arrowhead indicates newly detected possible intermediate compound with *m/z* 225 (without CoA). The vertical axis indicates intensity (count/sec) and the horizontal axis indicates RT (min).

**Fig 3.**
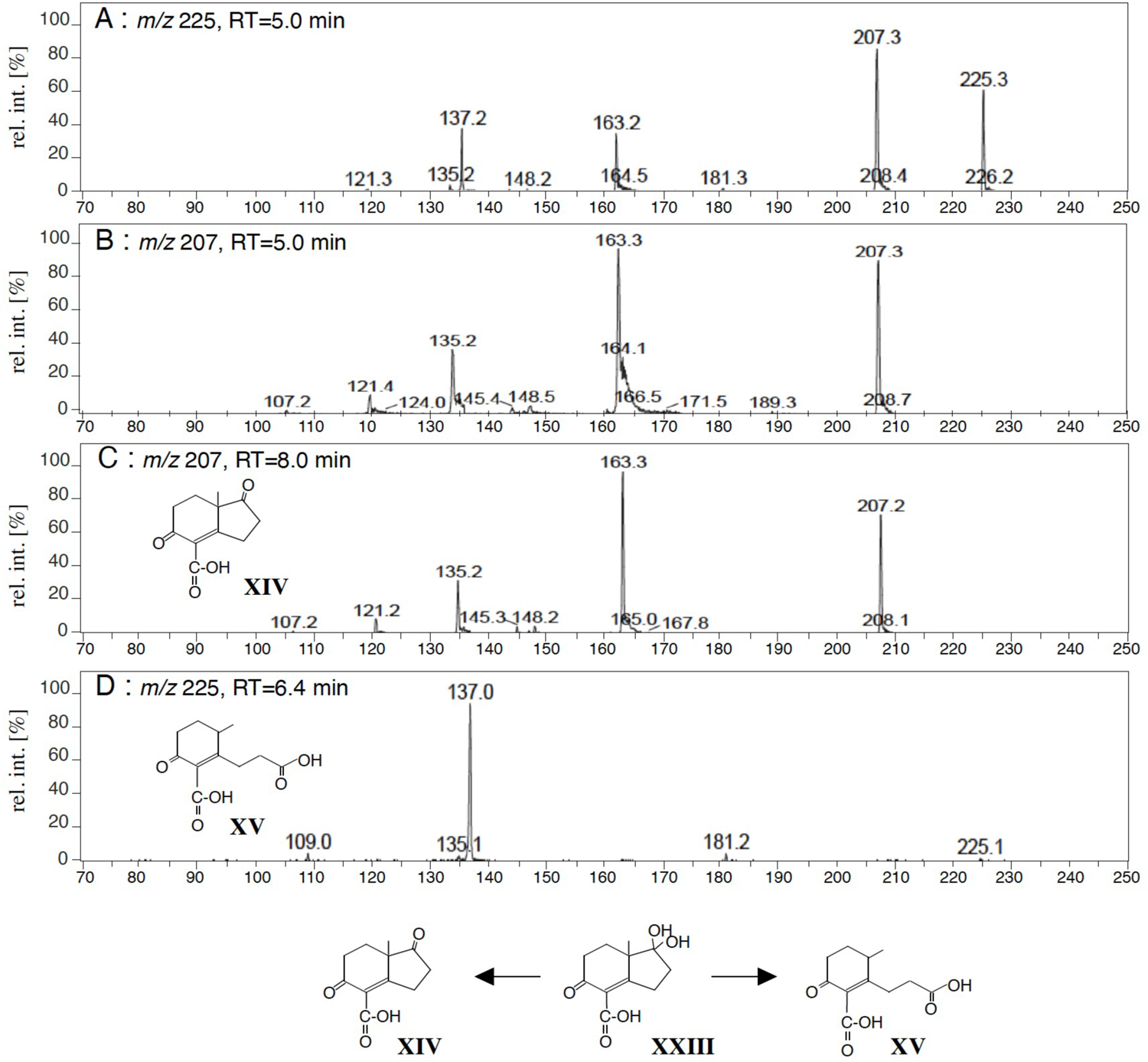
The mass spectra of a peak with *m/z* 225 at RT = 5.0 min. (A), a peak with *m/z* 207 at RT = 5.0 min (B), **XIV** (*m/z* 207, C), and **XV** (*m/z* 225, D) analyzed by LC/MS/MS. Compounds are; 9,17-dioxo-1,2,3,4,5,6,10,19-octanorandrost-8(14)-en-7-oic acid (**XIV**), 9-oxo-1,2,3,4,5,6,10,19-octanor-13,17-secoandrost-8(14)-ene-7,17-dioic acid (**XV**), and 17-dihydroxy-9-oxo-1,2,3,4,5,6,10,19-octanorandrost-8(14)-en-7-oic acid (**XXIII**). The vertical axis indicates relative intensity (%) and the horizontal axis indicates mass (*m/z*).

### Construction of plasmid pMFYMhpR for the induction of target genes in TA441 mutants

The degradation of C- and D-rings, as well as that of the cleaved B-ring proceeds via β-oxidation. The intermediates in this procedure are CoA-esters, which hinders their conversion using the purified enzyme. To investigate unknown intermediate compounds originating from C- and D-ring cleavage, we constructed a plasmid that enabled the induction of target genes in various TA441 mutants. Degradation of 3-(3-hydroxyphenyl)propionic acid (3HPP) by TA441 depends on induction of the Mhp gene cluster (39). Specifically, Mhp catalytic genes are induced following binding of 3HPP and the positive regulator MhpR to the promoter region (Fig. 4). The DNA fragment containing *mhpR* and the promoter region was amplified and cloned into the *Pvu*II site of the broad-host-range plasmid pMFY42 (40) to generate pMFYMhpR (Fig. 4).

**Fig 4.**
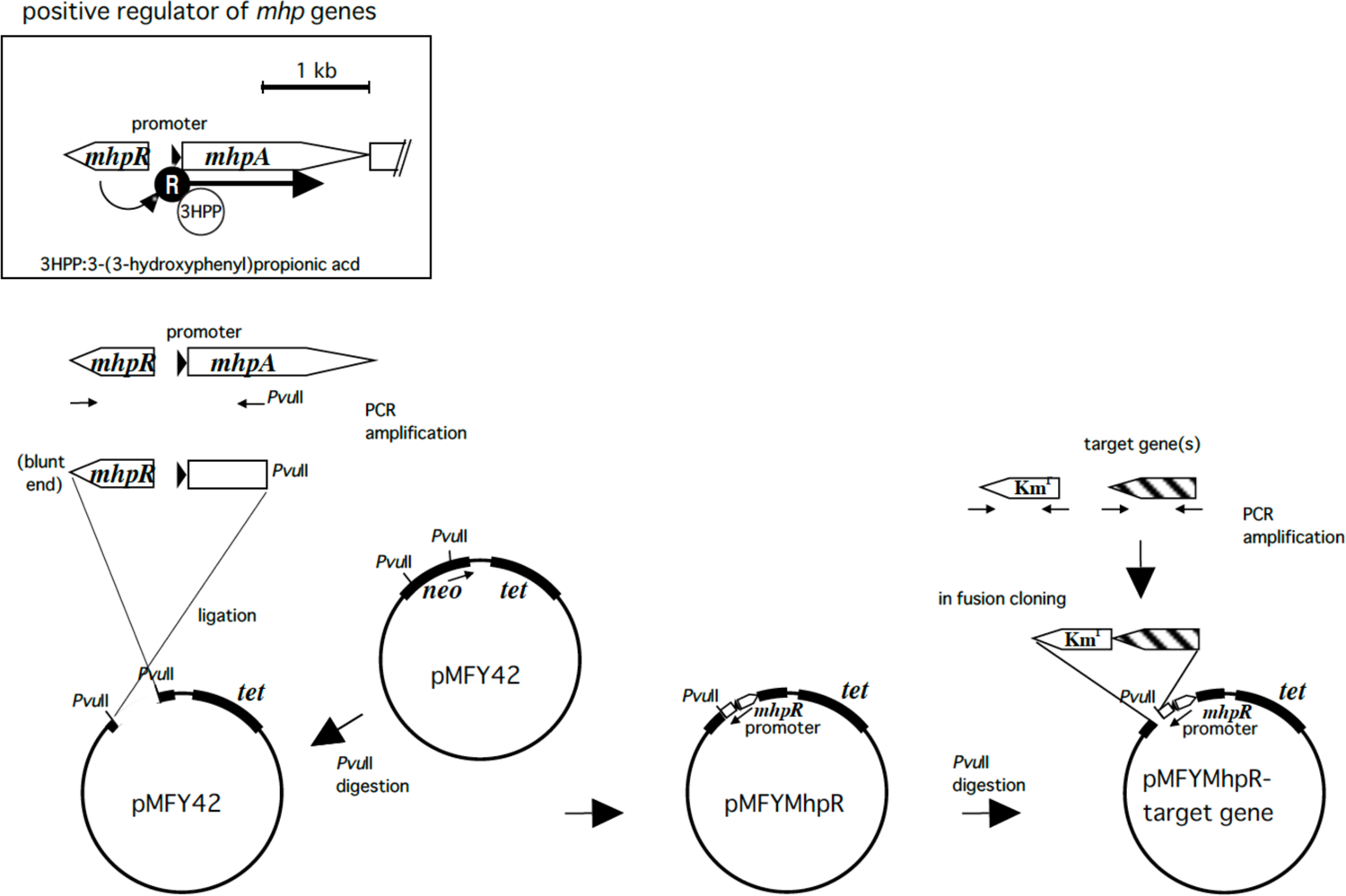
Construction of pMFYMhpR to induce objective gene(s) in TA441 mutants. TA441 degrades 3-(3-hydroxyphenyl)propionic acid (3HPP) with *mhpRABD* (42). MhpR is the positive regulator of the *mhp* genes. The conjugate of MhpR and 3HPP attached to the promotor region to express *mhpABD*. A DNA fragment containing *mhpR* and the promoter region was amplified using pYT11, a pUC19 derivative carrying *mhpRABD* (42), and cloned into the *Pvu*II site of the broad-host-range pMFY42 plasmid (43) using In-Fusion HD Cloning Kit (TAKARA Bio, Japan). Then PCR-amplified objective gene was cloned into the *Pvu*II site of pMFYMhpR with a kanamycin-resistance gene for selection. *tet*: tetracycline resistance gene. *neo*: kanamycin resistance gene.

To examine whether the obtained plasmid was indeed functional and whether **XXIII**-CoA ester was converted by ScdY, we introduced *scdY* into the *Pvu*II site downstream of the Mhp promoter on pMFYMhpR. The resulting pMFYMhpRScdY plasmid was transferred to ScdY^-^ mutants and these were incubated with cholic acid for 7 days. Then, 3HPP was added to the culture and the cells were incubated for another 3 days. Mass chromatograms of peaks with *m/z* 207 and *m/z* 181 (for detection of **XV’**, a decarboxylated derivative of **XV**) recorded at 0, 1, and 3 days after the addition of 3HPP are shown in Fig. 5. Chromatograms of peaks with *m/z* 225 (for detection of **XV**), *m/z* 183 (for detection of **XVII’**, a decarboxylated derivative of **XVII**), and *m/z* 197 (for detection of **XVI’**, a decarboxylated derivative of **XVI**) are shown in Fig. S1 in Supplemental Material. After 1 day, **XIV** and **XXIII** content was only slightly lower compared to that on day 0 (Fig. 5A and B), but it became barely detectable after 3 days (Fig. 5C). These results indicate that induction using the pMFYMhpR-based plasmid was successful and **XXIII**-CoA was the substrate of ScdY.

**Fig 5.**
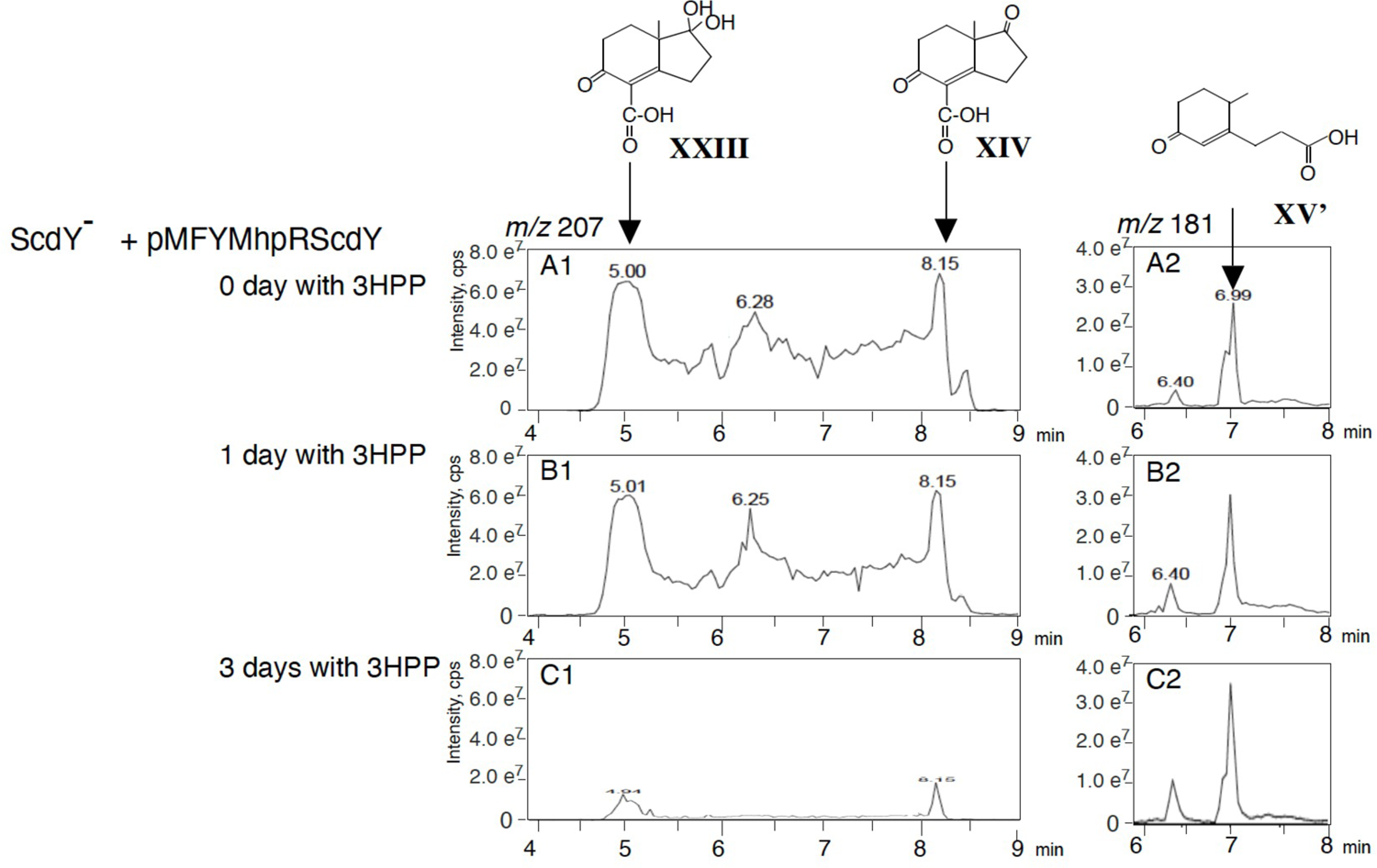
Induction of ScdY in the ScdY^-^ mutant and the conversion of **XXIII** by ScdY. ScdY^-^ carrying pMFYMhpRScdY was incubated with 0.1% cholic acid for 7 days, then 0.1% 3HPP was added for induction. Mass chromatograms of the culture (A1-C1, *m/z* 207; A2-C2, *m/z* 181) in 0 day (A1, A2), 1 day (B1, B2), and 3 days (C1, C2) after addition of 3HPP are shown. Compounds are; 9,17-dioxo-1,2,3,4,5,6,10,19-octanorandrost-8(14)-en-7-oic acid (**XIV**), 17-dihydroxy-9-oxo-1,2,3,4,5,6,10,19-octanorandrost-8(14)-en-7-oic acid (**XXIII**), and 9-oxo-1,2,3,4,5,6,10,19-octanor-13,17-secoandrost-8(14)-en-17-oic acid (**XV’**). The vertical axis indicates intensity (count/sec) and the horizontal axis indicates RT (min).

### Characterization of the product of ScdY

The product of ScdY and the derivatives detected in the previous section became only slightly more abundant after induction. This is because the product was further degraded by ScdY^-^ cells carrying pMFYMhpRScdY. Next, we constructed another mutant deficient for both ScdY and ScdL2. Because it was difficult to construct a mutant, in which only ScdL2 and ScdY were disrupted, ScdL2^-^N^-^Y^-^ was generated instead. ScdN acts downstream of ScdL1L2 and does not affect ScdY activity (Fig. 1). After pMFYMhpRScdY was introduced into ScdL2^-^N^-^Y^-^ cells, these were first incubated with cholic acid for 7 days and 3HPP for another 3 days. Samples were analyzed every 24 h using LC-MS/MS. Mass chromatograms of peaks with *m/z* 181 (**XV’**), *m/z* 183 (**XVII’**), *m/z* 197 (**XVI’**), and *m/z* 229 (9-hydroxy-1,2,3,4,5,6,10,19-octanor-13,17-secoandrostane-7,17-dioic acid [**XXIV**]) are shown in Fig. 6. **XV’**, **XVII’**, **XVI’**, and **XXIV** content increased with induction time. The increase appeared slightly slower for **XVII’** and **XXIV**, which have a hydroxyl group at C9, compared to **XV’** and **XVI**, which have a ketone moiety at C9. In addition to these four compounds, which were always found in ScdL1^-^L2^-^ cultures, a compound with *m/z* 181 at RT = 7.4 min was detected (**XXV’**). The amount of **XXV’** increased during the first 2 days, but started to decrease thereafter, which was confirmed by various repetitions. When re-examining past data, it emerged that **XXV’** was occasionally detected at low amounts in the ScdL1^-^L2^-^ culture (for example, Supplemental Material Fig. S1 in reference [36]) and its mass spectrum was almost identical to that of **XV’** (Fig. 6D). In our experiments, compounds with a C9 ketone and C8(14) single bond, as well as those with a C9 hydroxyl group and C8(14) double bond were unstable and were converted to compounds with a C9 ketone and C8(14) double bond or those with a C9 hydroxyl group and C8(14) single bond after addition of CoA. Based on the given evidence, the unknown compound with *m/z* 181 at RT = 7.4 min was suggested to be a decarboxylated derivative of 9-oxi-1,2,3,4,5,6,10,19-octanor-13,17-secoandrost-13(14)-ene-7,17-dioic acid (**XXV**) (Fig. 6D2). Even though ScdL2^-^N^-^Y^-^ cells carrying pMFYMhpRScdY accumulated substantial amounts of **XVI**, its role remains unclear. **XVI** was detected in ScdL1^-^L2^-^ cultures incubated with a steroid compound containing or not a C12-hydroxyl group.

**Fig 6.**
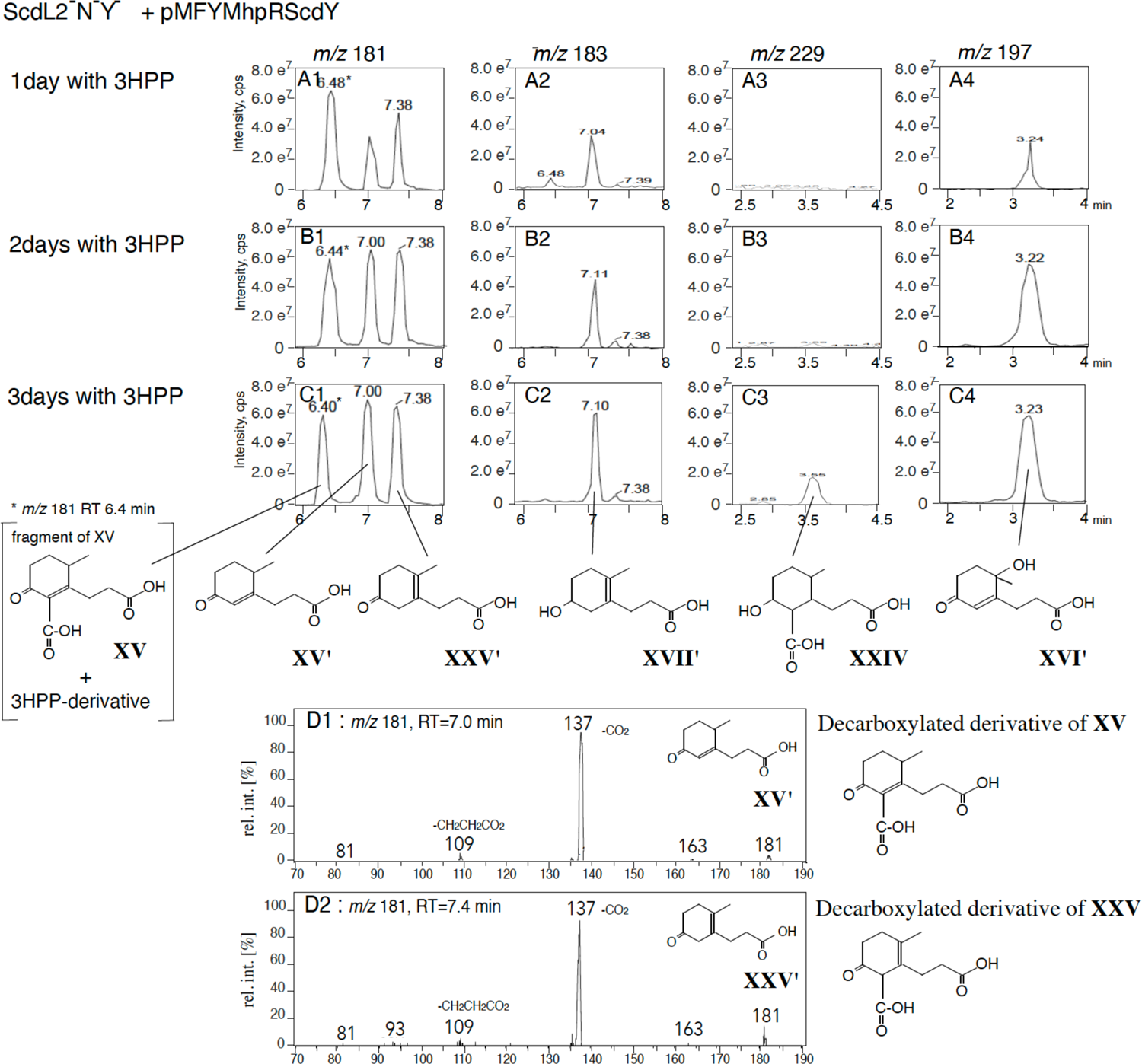
Induction of ScdY in the ScdL2^-^N^-^Y^-^ mutant (ScdN acts downstream of ScdL1L2 and does not affect ScdY activity). ScdL2^-^N^-^Y^-^ carrying pMFYMhpRScdY was incubated with 0.1% cholic acid for 7 days, then 0.1% 3HPP was added for induction. Mass chromatograms (A1-C1, *m/z* 181; A2-C2, *m/z* 183; A3-C3, *m/z* 229; A4-C4, *m/z* 197) of 1 day (A), 2 days (B), and 3 days (C) after addition of 3HPP are shown. D1 and D2 are the mass spectrum of **XV’** and a peak with *m/z* 181 at RT = 7.4 min, respectively. *On the mass chromatogram of *m/z* 181, both a fragment of **XV** and a derivative of 3HPP are detected exactly at the same RT of 6.4 min. Compounds are; 9-oxo-1,2,3,4,5,6,10,19-octanor-13,17-secoandrost-8(14)-ene-7,17-dioic acid (**XV**), 9-oxo-1,2,3,4,5,6,10,19-octanor-13,17-secoandrost-8(14)-en-17-oic acid (**XV’**), 9-oxo-1,2,3,4,5,6,10,19-octanor-13,17-secoandrost-13-en-17-oic acid (**XXV’**), 9-hydroxy-1,2,3,4,5,6,10,19-octanor-13,17-secoandrost-13-en-17-oic acid (**XVII’**), 9-hydroxy-1,2,3,4,5,6,10,19-octanor-13,17-secoandrostane-7,17-dioic acid (**XXIV**), and 13-hydroxy-9-oxo-1,2,3,4,5,6,10,19-octanor-13,17-secoandrost-8(14)-en-17-oic acid (**XVI’**). In mass chromatograms, the vertical axis indicates intensity (count/sec) and the horizontal axis indicates RT (min) (A-C). In the mass spectrum, the vertical axis indicates relative intensity (%) and the horizontal axis indicates mass (*m/z*) (D).

### Characterization of the product of ScdL1L2

4-Methyl-5-oxo-octane-1,8-dioic acid (**XX**)-CoA ester is the major substrate of ScdM1M2 (Fig. 1), and a large amount of **XX** accumulates in ScdM1^-^M2^-^ cultures (36). **XX** is generated from the product of C-ring cleavage by the CoA transferase ScdJ. We expected this product to be **XIX**-CoA ester (MW 244) (Fig. 1), however, a compound with *m/z*243 was not detected in the mass chromatogram of the ScdJ^-^ culture (see Fig. S3A1 in Supplemental Material). Then the *scdL1L2* gene was introduced downstream of the Mhp promoter of pMFYMhpR, and the resulting pMFYMhpRScdL1L2 plasmid was transformed into ScdL1^-^L2^-^ cells, which were incubated first with cholic acid for 7 days and then 3HPP for another 3 days. No suitable products or derivatives were detected (data not shown). When 3HPP and cholic acid were added at the beginning of the incubation and the cells were cultured for 7 days, peaks corresponding to possible ScdL1L2 products (*m/z* 245 at RT = 5.6 min, *m/z* 241 at RT = 5.7 min, and *m/z* 227 at RT = 7.0 min and 7.7 min) were detected (Fig. 7A1–3 and 7B1–3). Mass chromatograms of peaks with *m/z* 243, 245, 241, 227, 225, and 229 from RT = 0 min to RT = 12 min are shown in Fig. S2 in Supplemental Material. A peak with *m/z* 241 at RT = 5.7 min was not detected when the initial steroid did not contain a C12-hydroxyl group. To investigate the changes in the amount of compounds induced by ScdL1L2, we constructed another plasmid, pMFYMhpRScdL1L2Y, carrying *scdL1L2Y*, and introduced it into ScdL2^-^N^-^Y^-^ cells. ScdJ is the main CoA transferase for the ring cleavage product of ScdL1L2. A double mutant of ScdL1L2 and ScdJ was the most suitable candidate in which to assess ScdL1L2 activity, but the 10-kb distance between *scdJ* and *scdL1L2* hindered construction of the mutant. We expected ScdL2^-^N^-^Y^-^ cells carrying pMFYMhpRScdL1L2Y to accumulate more putative ScdL1L2 products following addition of 3HPP because ScdN catalyzes two steps downstream of C-ring cleavage (Fig. 1). ScdL2^-^N^-^Y^-^ cells carrying pMFYMhpRScdL1L2Y were incubated with cholic acid for 7 days and 3HPP for another 3 days. Induction of ScdL1L2 led to the accumulation of the four candidate compounds (Fig. 7D), particularly the one with *m/z* 227 (Fig. 7D3). These results suggested that these four compounds were the products of ScdL1L2 or their derivatives. To boost their accumulation, we constructed one more mutant, ScdJ^-^, which did not accumulate any compounds with MW 244, but presented large amounts of **XV’**, **XVII’**, and **XVI’** (36). ScdJ^-^ carrying pMFYMhpRScdL1L2 was expected to accumulate more of these compounds following 3HPP induction. LC-MS/MS analysis of ScdJ^-^ carrying pMFYMhpRScdL1L2 revealed a drastic increase of the peak with *m/z* 245 at RT = 5.6 min and peaks with *m/z* 227 at RT = 7.0 min and 7.7 min in the induced cultures (Fig. 7E–G). The mass spectrum of the peak with *m/z* 245 at RT = 5.6 min indicated that the compound was 3-hydroxy-6-methyl-7-oxo-decane-1,10-dioic acid (**XXVI**) (Fig. 8A). The peaks with *m/z* 227 at RT = 7.0 min and 7.7 min were identified as the dehydrated derivatives of **XXVI**, 6-methyl-7-oxo-dec-3-ene-1,10-dioic acid (**XXVII**) and 6-methyl-7-oxo-dec-2-ene-1,10-dioic acid (**XXVIII**), respectively (Fig. 8B and C). The mass spectrum of the peak with *m/z* 241 at RT = 5.7 min assigned the compound to 6-methyl-3,7-dioxo-dec-5-ene-1,10-dioic acid (**XXIX**) (Fig. 8D). Finally, **XXIX** was thought to be a dehydrated derivative of 5-hydroxy-6-methyl-3,7-dioxo-decane-1,10-dioic acid (**XXX**). These results indicate that **XXVI**-CoA ester is the major product of C-ring cleavage by ScdL1L2. During β-oxidation, **XVIII**-CoA ester is produced from **XXVI**-CoA ester. Here, **XVIII** was almost undetectable, probably because it was readily converted to **XX**-CoA ester by ScdJ or other CoA transferases such as ScdF, allowing only a small amount of **XX** to be detected in the ScdJ^-^ culture (36). The peak with *m/z* 245 at RT = 1.7 min detected in ScdL1^-^L2^-^ cells carrying pMFYMhpR was hypothesized to be **XXXI**. This is because the overall fragment pattern of the mass spectrum was similar to that of **XVIII**, and fragments of *m/z* 123 and *m/z* 129, which are characteristic of **XXVI**, were detected in the mass spectrum of this peak (see Fig. S4 in Supplemental Material). The results also show that C- and D-ring cleavage proceeded in the presence of the C12-hydroxyl group, but degradation was less efficient than for compounds devoid of the C12-hydroxyl group.

**Fig 7.**
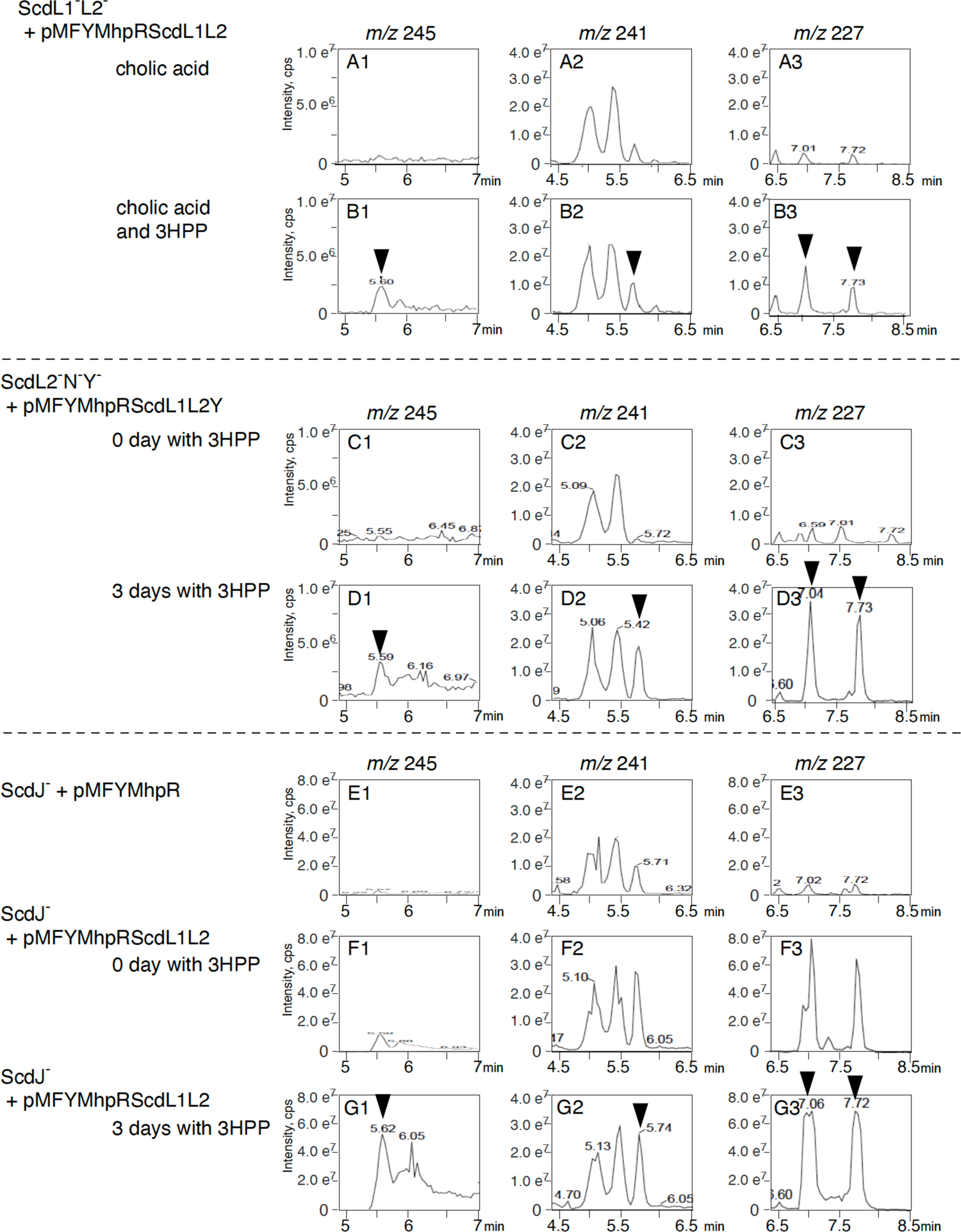
Induction of ScdL1L2 in the ScdL1^-^L2^-^ mutant. (A,B), ScdL1L2Y in the ScdL2^-^N^-^Y^-^ mutant (C,D), and ScdL1L2 in the ScdJ^-^ mutant (F,G). ScdL1^-^L2^-^ carrying pMFYMhpRScdL1L2 was incubated with 0.1% cholic acid (A) and with 0.1% cholic acid and 0.1% 3HPP (B) for 7 days. ScdL2^-^N^-^Y^-^ carrying pMFYMhpRScdL1L2Yc(C, D), ScdJ^-^ carrying pMFY42 (E, negative control), and ScdJ^-^ carrying pMFYMhpRScdL1L2 (F, G) were incubated with 0.1% cholic acid for 7days. Then 0.1% 3HPP was added for induction and incubated for another 3 days (D,G) (ScdJ^-^ carrying pMFY42 incubated for 3 days with 3HPP is not shown since it was almost the same as E). Arrowheads indicate possible product of ScdL1L2 and the derivatives. Mass chromatograms of *m/z* 245 (A1-G1), *m/z* 241 (A2-G2) and *m/z* 227 (A3-G3) are shown. The vertical axis indicates intensity (count/sec) and the horizontal axis indicates RT (min).

**Fig 8.**
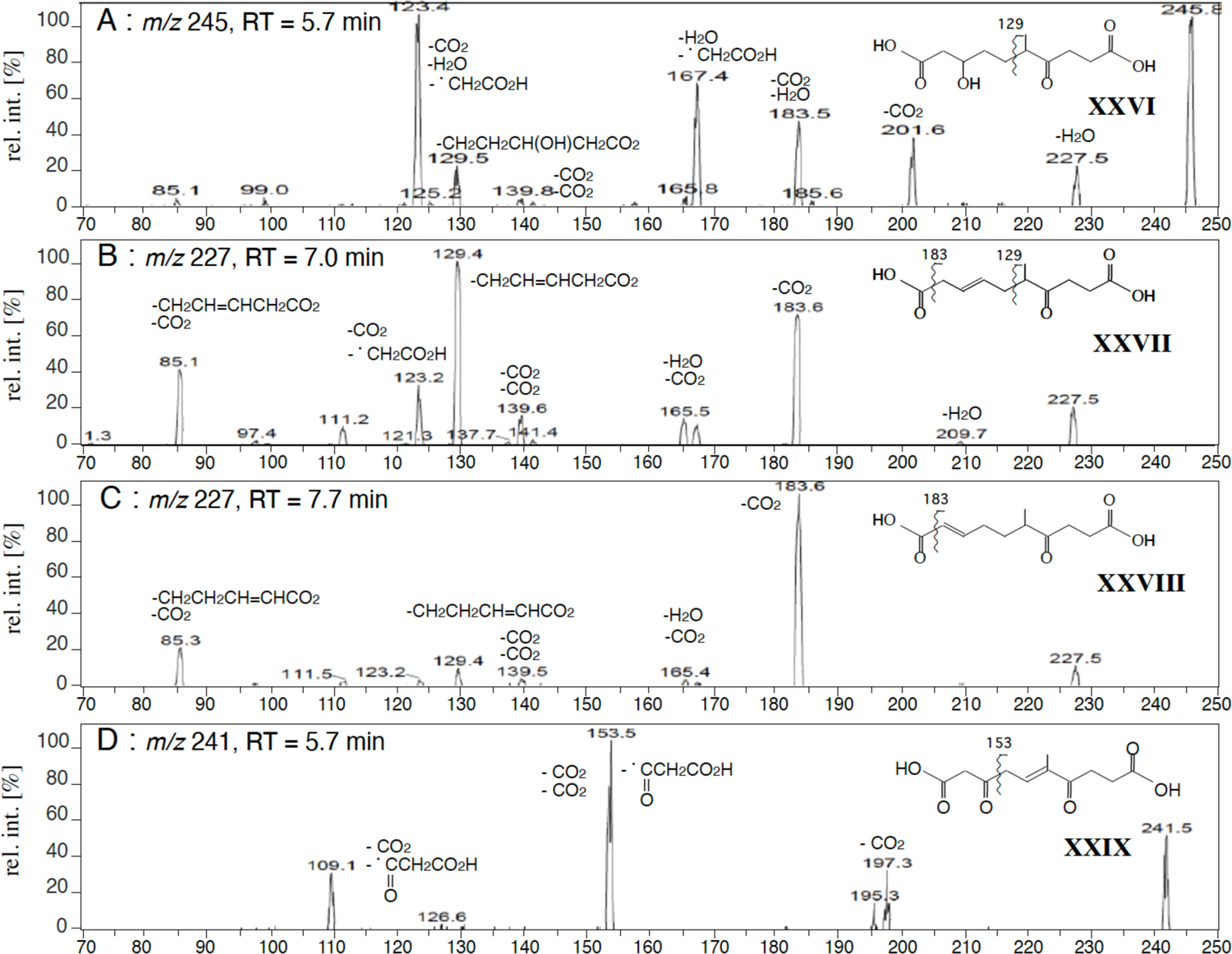
The mass spectra of a peak with *m/z* 245 at RT = 5.6 min. (A), a peak with *m/z* 227 at RT = 7.0 min (B), a peak with *m/z* 227 at RT = 7.7 min (C), and a peak with *m/z* 241 at RT = 5.7 min (D). Compounds are; 3-hydroxy-6-methyl-7-oxo-decane-1,10-dioic acid (**XXVI**), 6-methyl-7-oxo-dec-3-ene-1,10-dioic acid (**XXVII**), 6-methyl-7-oxo-dec-2-ene-1,10-dioic acid (**XXVIII**), and 6-methyl-3,7-dioxo-dec-5-ene-1,10-dioic acid (**XXIX**). The vertical axis indicates relative intensity (%) and the horizontal axis indicates mass (*m/z*).

## Conclusion

The degradation pathway emerging from this study is illustrated in Fig. 9. The substrate of the ScdY hydratase is a geminal diol, **XXIII**-CoA ester. Hydration at C14 by ScdY leads to cleavage of the D-ring at C13-C17 and the product, **XVIII**-CoA ester, is likely hydrogenated at C9. This finding is supported by **XXVI**-CoA ester being the major C-ring cleaved compound produced by ScdL1L2, and by accumulation of **XVII’** and **XXIV**, both of which have a C9 hydroxyl group, in ScdL1^-^L2^-^ cultures. C- and D-ring cleavage can proceed in the presence of the C12-hydroxyl group; however, the main degradation pathway relies on removal of the C12-hydroxyl group before addition of CoA by ScdA. Accumulating data revealing complex interplay between bile acids, gut microbiota, and host metabolism have shed new light on a potential impact of bile acids on the brain-gut-microbiota axis (39–41). Precise bacterial bile acid degradation pathway will be one of the key to investigate the gut–microbiota–brain axis.

**Fig 9.**
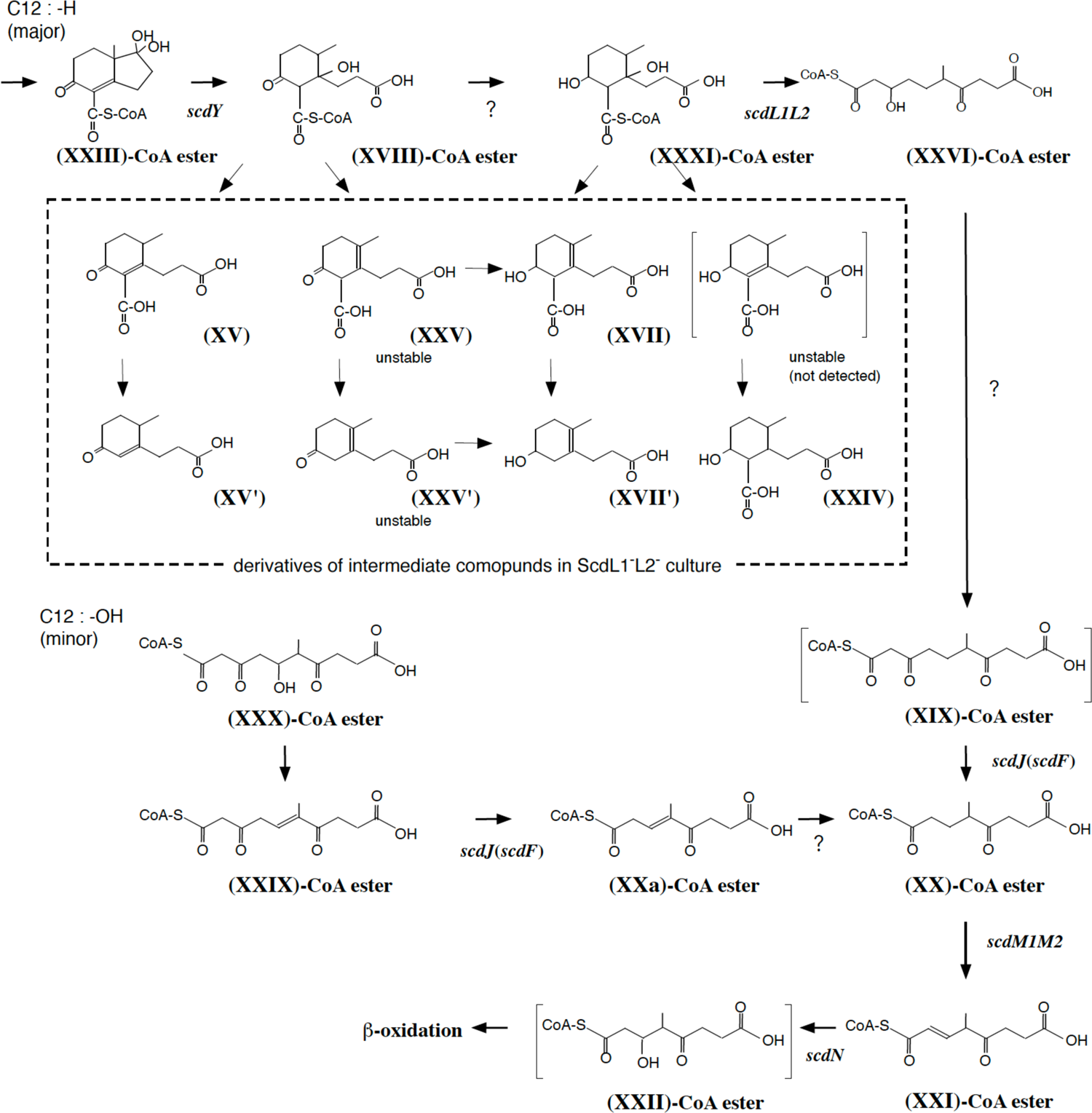
Proposed C- and D-ring cleavage process. Compounds in the broken square are derivatives identified in the culture of ScdL1^-^L2^-^ mutant in the previous study. Compounds in brackets are speculation. Compounds are; 17-dihydroxy-9-oxo-1,2,3,4,5,6,10,19-octanorandrost-8(14)-en-7-oic acid (**XXIII**), 14-hydroxy-9-oxo-1,2,3,4,5,6,10,19-octanor-13,17-secoandrostane-7,17-dioic acid (**XVIII**), 9,14-dihydroxy-1,2,3,4,5,6,10,19-octanor-13,17-secoandrostane-7,17-dioic acid (**XXXI**), 3-hydroxy-6-methyl-7-oxo-decane-1,10-dioic acid (**XXVI**), 6-methyl-3,7-dioxo-decane-1,10-dioic acid (**XIX**), 4-methyl-5-oxo-octane-1,8-dioic acid (**XX**), 4-methyl-5-oxo-oct-2-ene-1,8-dioic acid (**XXI**), 3-hydroxy-4-methyl-5-oxo-octane-1,8-dioic acid (**XXII**), 5-hydroxy-6-methyl-3,7-dioxo-decane-1,10-dioic acid (**XXX**), 6-methyl-3,7-dioxo-dec-5-ene-1,10-dioic acid (**XXIX**), 4-methyl-5-oxo-oct-3-ene-1,8-dioic acid (**XXa**), 9-oxo-1,2,3,4,5,6,10,19-octanor-13,17-secoandrost-8(14)-ene-7,17-dioic acid (**XV**), 9-oxo-1,2,3,4,5,6,10,19-octanor-13,17-secoandrost-8(14)-en-17-oic acid (**XV’**), 9-oxo-1,2,3,4,5,6,10,19-octanor-13,17-secoandrost-13-ene-7,17-dioic acid (**XXV**), 9-oxo-1,2,3,4,5,6,10,19-octanor-13,17-secoandrost-13-en-17-oic acid (**XXV’**), 9-hydroxy-1,2,3,4,5,6,10,19-octanor-13,17-secoandrost-13-ene-7,17-dioic acid (**XVII**), 9-hydroxy-1,2,3,4,5,6,10,19-octanor-13,17-secoandrost-13-en-17-oic acid (**XVII’**), 9-hydroxy-1,2,3,4,5,6,10,19-octanor-13,17-secoandrostane-7,17-dioic acid (**XXIV**), and 9-hydroxy-1,2,3,4,5,6,10,19-octanor-13,17-secoandrost-8(14)-ene-7,17-dioic acid (**XXXI**). Mass spectrum of the candidate of **XXXI** (a peak of *m/z* 245 at RT = 1.7 min and 2.75 min) is presented in the supplementary material Fig. S4.

## MATERIALS AND METHODS

### Abbreviations

LC/MS/MS, reverse-phase liquid chromatography with tandem mass spectrometry; RT, retention time; MW, molecular weight; CoA, coenzyme A; 3HPP, 3-(3-Hydroxyphenyl)propionic acid; PCR, polymerase chain reaction.

### Culture conditions

Mutant strains of *C. testosteroni* TA441 were grown at 30°C in a mixture of equal volumes of Luria-Bertani (LB) medium and C medium (a mineral medium for TA441) (22) with suitable carbon sources. This mixed media is used because the mutants accumulate more amount of intermediate compounds than with C medium or LB medium (unpublished data). Cholic acid and other steroids were added as filter-sterilized DMSO solutions with a final concentration of 0.1% (w/v). 3-(3-Hydroxyphenyl)propionic acid (3HPP) was added as acetonitrile solution with a final concentration of 0.1% (w/v).

### Construction of deletion mutants, plasmids, and mutants for complementation experiments

To construct the ScdL2^-^N^-^Y^-^ mutant (Table 1), a DNA fragment containing *scdK* and *scdY*::Km^r^ (bearing the kanamycin resistance gene) was amplified using the pUC19-based plasmid pUCORF5(*scdY*)-Km^r^, which had been generated in a previous study (34) to delete *scdY*. A DNA fragment containing *scdM1M2F* was amplified using pUCScdM1M2F, a pUC19 derivative (Tables 2 and 3). Then, the two fragments were introduced into a pHSG397 derivative carrying *tesB* and *scdL1* using the In-Fusion HD Cloning Kit (TaKaRa Bio, Japan). This yielded the pHSGScdL2NY-Km^r^ plasmid carrying *tesB*, *scdL1*, *scdK*, *scdY*::Km^r^, and *scdM1M2F.* The plasmid was introduced into TA441 cells via electroporation. Successful transformants were selected on LB plates with kanamycin and chloramphenicol. Deletion of *scdL2*, insertion of Km^r^ in *scdY,* and the presence of *scdK* were confirmed by PCR amplification. To construct pMFYMhpR, a DNA fragment containing *mhpR* and the promoter region was amplified using pYT11, a pUC19 derivative carrying *mhpRABD* (42), and cloned into the *Pvu*II site of the broad-host-range pMFY42 plasmid (Fig. 4) (43). pMFYMhpR contained a unique *Pvu*II site downstream of the promoter. The target genes were PCR-amplified with Km^r^ for subsequent selection and introduced into the *Pvu*II site of pMFYMhpR using the In-Fusion HD Cloning Kit.

**Table 1.**
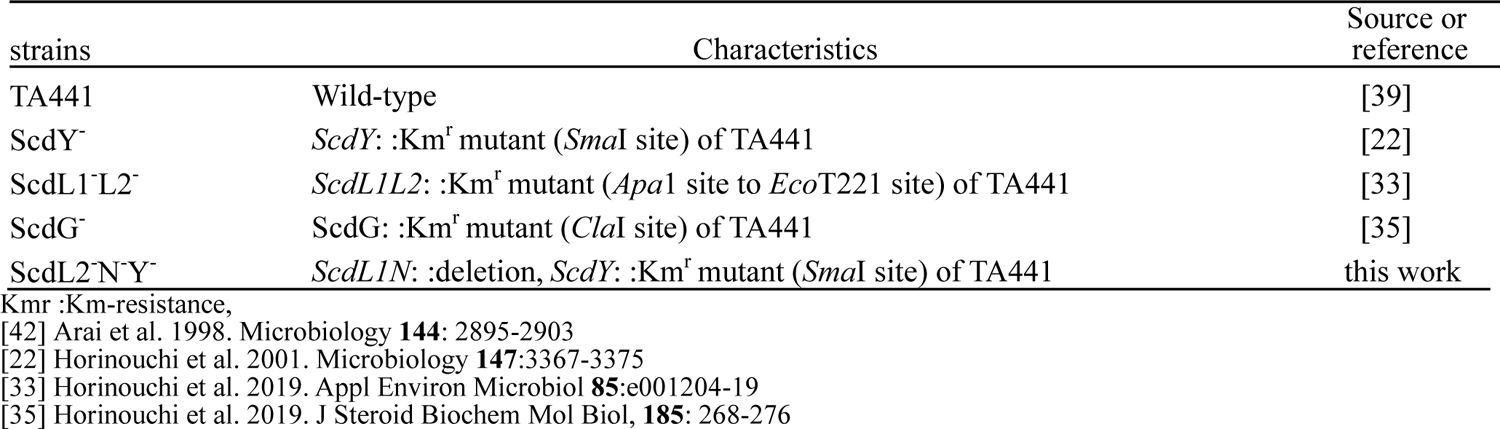
strains

**Table 2.** plasmids

| plasmids | Characteristics | Source or reference |
| --- | --- | --- |
| pUC19 | Ap<sup>r</sup>, *lacZ* | [41] |
| pMFY42 | Tc<sup>r</sup>, Km<sup>r</sup>, RSF1010-based broad host range plasmid | [40] |
| pYT11 | pUC19 derivative carrying *mhpRABD* (5.2 kb *Bgl*II-fragment) | [39] |
| pMFYMhpRA | pUC19 derivative carrying *mhpR* and the promoter of *mhp* genes | this work |
| pMFYMhpScdY | pMFYMhpRA derivative carrying *scdY* | this work |
| pMFYMhpScdL1L2 | pMFYMhpRA derivative carrying *scdL1L2* | this work |
| pMFYMhpScdL1L2Y | pMFYMhpRA derivative carrying *scdL1L2Y* | this work |
| pUCORF5(*scdY*)-Km<sup>r</sup> | pUC19 derivative carrying DNA fragment containing *scdK* and *scdY*: :Km<sup>r</sup> (*Sma*I site) | [34] |
| pHSGtesBscdL1 | pHSG397 derivative carrying *tesB* and *ScdL1* | this work |
| pUCScdM1M2F | pUC19 derivative carrying *scdM1M2F* | this work |
| pHSGScdL2NY-Km<sup>r</sup> | pHSGtesBscdL1 derivative with *scdK*, *scdY*: :Km<sup>r</sup>, and *scdM1M2F* | this work |

Km<sup>r</sup>: Km-resistance pSuperCos1\* (Stratagene, CA) [44] Vieira, J., and Messing, J. 1987. Methods Enzymol. **153**: 3-11. [43] Nagata Y. et al. 1993. J Bacteriol **175**:6403-6410 [42] Arai H. et al. 1999. Microbiology **145**:2813-2820. [34] Horinouchi M, et al. 2018. Appl Environ Microbiol **84**:e01324-18.

**Table 3.**
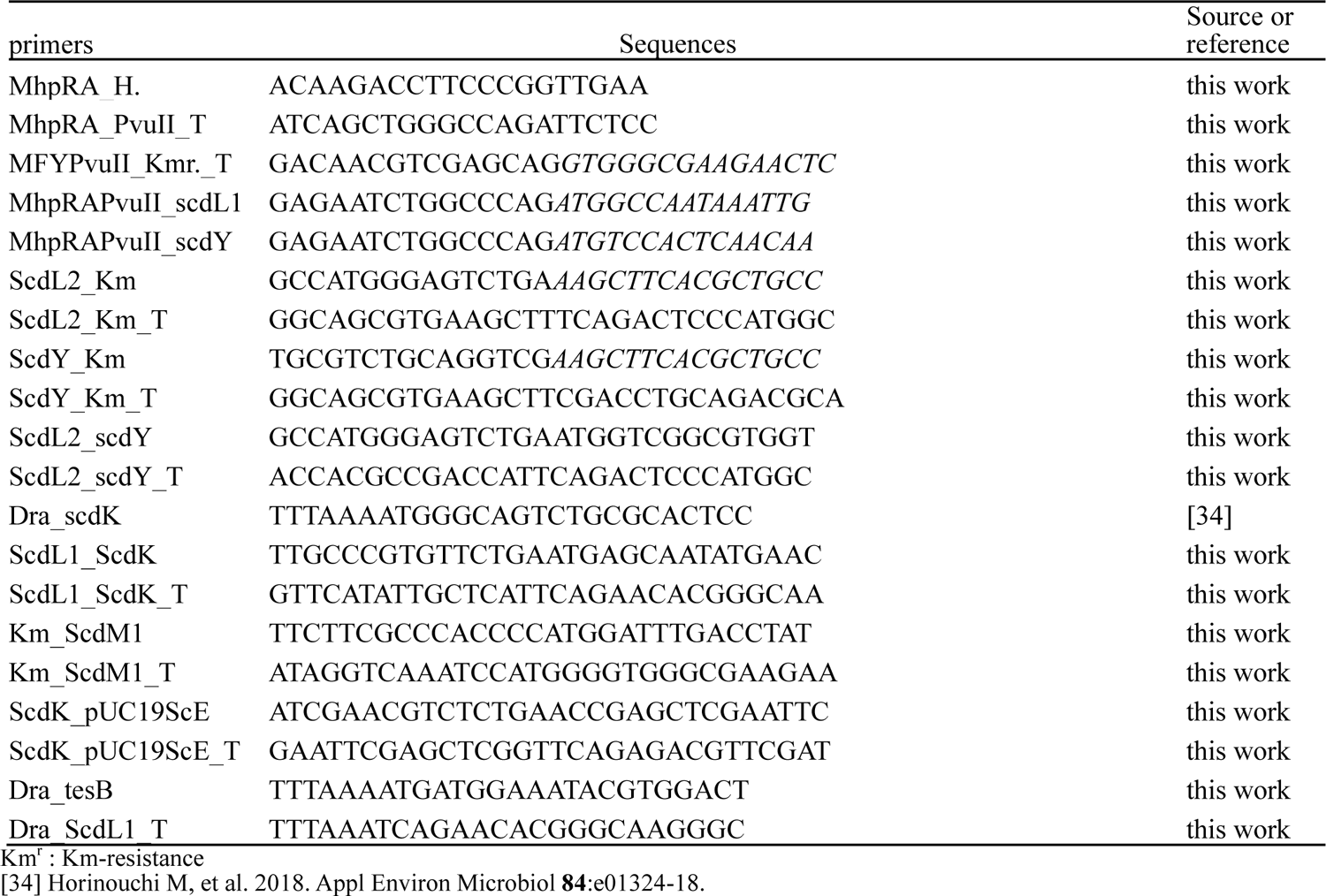
primers

### Reverse-phase liquid chromatography with tandem mass spectrometry (LC/MS/MS)

For LC/MS/MS analysis, 2 µl of the samples prepared the same way as those for HPLC/MS analysis was injected into the system. Agilent 1100 HPLC (Agilent, CA) with a mass spectrometer (Applied Biosystems 4000 Q-TRAP, MS) was used with L-column2 ODS (1.5 × 150 mm) Type L2-C 18.5µm, 12mm (GL Science, Tokyo, Japan) and elution was carried out using 90% solution C (H_2_O:HCOOH = 100:0.1) and 10% solution A for 1min, followed by a linear gradient from 90% solution C and 10% solution A to 20% solution C and 80% solution A over 7 min, which was maintained for 2 min. The flow rate was 0.2 ml/min. The desolvation temperature for the mass spectrometer was fixed at 450 °C. The collision energy was 20 V.

### Sample preparation for LC/MS/MS

500 μl of the culture was acidified with HCl (pH2) and extracted with 1ml ethyl acetate twice. The ethyl acetate layer was dried, resolved in 600 μl methanol, stored at −60°C, and 2 μl of the methanol solution was subjected to LC/MS/MS analysis within 2d

### Data availability

The authors agree that any materials and data that are reasonably requested by others are available from a publicly accessible collection or will be made available in a timely fashion, at reasonable cost, and in limited quantities to members of the scientific community for noncommercial purposes.

## ACKNOWLEDGMENTS

MH appreciates Dr. Reizo Kato (head of Condensed Molecular Materials Laboratory, RIKEN) and Dr. Yousoo Kim (head of Surface and Interface Science Laboratory, RIKEN) for thoughtful supports and advice. The authors thank Dr. Takemichi Nakamura (Molecular Structure Characterization Unit, RIKEN CSRS, WAKO) for his assistance in mass spectrometry and Dr. Hiroyuki Koshino (Molecular Structure Characterization Unit, RIKEN CSRS, WAKO) for his assistance in NMR analysis.

## FIGURE LEGENDS (SUPPLEMENTARY MATERIAL)

Fig S1 Additional data for Fig. 5. Induction of ScdY in the ScdY^-^ mutant. ScdY^-^ carrying pMFYMhpRScdY was incubated with 0.1% cholic acid for 7days and 0.1% 3HPP was added for induction. Mass chromatograms of *m/z* 207 (A1-C1), *m/z* 225 (A2-C2), *m/z* 181 (A3-C3, *m/z* 183 (A4-C4), and *m/z* 197 (A5-C5) in 0 day (A1-5), 1 day (B1-5), and 3 days (C1-5) after addition of 3HPP are shown. Compounds are; 9,17-dioxo-1,2,3,4,5,6,10,19-octanorandrost-8(14)-en-7-oic acid (**XIV**), 17-dihydroxy-9-oxo-1,2,3,4,5,6,10,19-octanorandrost-8(14)-en-7-oic acid (**XXIII**), 9-oxo-1,2,3,4,5,6,10,19-octanor-13,17-secoandrost-8(14)-en-17-oic acid (**XV’**), 9-hydroxy-1,2,3,4,5,6,10,19-octanor-13,17-secoandrost-13-en-17-oic acid (**XVII’**), and 13-hydroxy-9-oxo-1,2,3,4,5,6,10,19-octanor-13,17-secoandrost-8(14)-en-17-oic acid (**XVI’**). The vertical axis indicates intensity (count/sec) and the horizontal axis indicates RT (min).

Fig S2 Additional data for Fig. 7A and 7B. Induction of ScdL1L2 in the ScdL1^-^L2^-^ mutant. ScdL1^-^L2^-^ carrying pMFYMhpRScdL1L2 incubated with 0.1% cholic acid for 7 days (A1-F1) and incubated with 0.1% cholic acid and 0.1% 3HPP for 7days (A2-F2). Chromatograms of *m/z* 243 (A), *m/z* 245 (B), *m/z* 241 (C), *m/z* 227 (D), *m/z* 225 (E), and *m/z* 229 (F) from RT = 0 min to 12 min are shown. Open arrowheads indicate peaks reduced in the culture with induction and closed arrowheads indicate peaks increased in the culture with induction. The vertical axis indicates intensity (count/sec) and the horizontal axis indicates RT (min). Compounds are; 14-hydroxy-9-oxo-1,2,3,4,5,6,10,19-octanor-13,17-secoandrostane-7,17-dioic acid (**XVIII**), 9,14-dihydroxy-1,2,3,4,5,6,10,19-octanor-13,17-secoandrostane-7,17-dioic acid (**XXXI**), 3-hydroxy-6-methyl-7-oxo-decane-1,10-dioic acid (**XXVI**), 13-hydroxy-9-oxo-1,2,3,4,5,6,10,19-octanor-13,17-secoandrost-8(14)-ene-7,17-dioic acid (**XVI**), 6-methyl-3,7-dioxo-dec-5-ene-1,10-dioic acid (**XXIX**), 5-hydroxy-6-methyl-3,7-dioxo-decane-1,10-dioic acid (**XXX**), 9-hydroxy-1,2,3,4,5,6,10,19-octanor-13,17-secoandrost-13-ene-7,17-dioic acid (**XVII**), 6-methyl-7-oxo-dec-3-ene-1,10-dioic acid (**XXVII**), 6-methyl-7-oxo-dec-2-ene-1,10-dioic acid (**XXVIII**), 9-oxo-1,2,3,4,5,6,10,19-octanor-13,17-secoandrost-8(14)-ene-7,17-dioic acid (**XV**), and 9-hydroxy-1,2,3,4,5,6,10,19-octanor-13,17-secoandrostane-7,17-dioic acid (**XXIV**).

Fig S3 Additional data for Fig. 7E, 7F, and 7G. Induction of ScdL1L2 in the ScdJ^-^ mutant. ScdJ^-^ carrying pMFY42 was incubated with 0.1% cholic acid for 7 days (A1-C1, negative control), and ScdJ^-^ carrying pMFYMhpRScdL1L2 was incubated with 0.1% cholic acid for 7 days (A2-C2), then 0.1% 3HPP was added and incubated for 3 days (A3-C3). Chromatograms of *m/z* 243 (A), *m/z* 245 (B), and *m/z* 241 (C) from RT = 0 min to 12 min are shown. Arrowheads indicate peaks of the compounds on the right side of the columns. The vertical axis indicates intensity (count/sec) and the horizontal axis indicates RT (min). Compounds are; 6-methyl-3,7-dioxo-decane-1,10-dioic acid (**XIX**), 3-hydroxy-6-methyl-7-oxo-dec-5-ene-1,10-dioic acid (**XXXII**), 3,5-dihydroxy-6-methyl-7-oxo-decane-1,10-dioic acid (**XXXIII**), 3-hydroxy-6-methyl-7-oxo-decane-1,10-dioic acid (**XXVI**), 6-methyl-3,7-dioxo-dec-5-ene-1,10-dioic acid (**XXIX**), and 5-hydroxy-6-methyl-3,7-dioxo-decane-1,10-dioic acid (**XXX**).

Fig S4 Mass chromatograms (A) and mass spectrum (C) of the candidate for **XXXI**. Data from the same experiment as Fig. S2. Mass spectra of the peaks with *m/z* 243 at RT = 1.8 min and 2.8 min (A1), and peaks with *m/z* 245 at RT = 1.75 min and 2.75 min (A3) were almost identical, respectively. B, C, and D show the mass spectrum of **XVIII**, the peak with *m/z* 245 at RT = 1.75 min, and **XXVI**, respectively. Corresponding fragments in B and C are connected with lines. Characteristic fragments found both in the mass spectra of the peak with *m/z* 245 at RT = 1.75 min and **XXVI** are indicated with closed arrowheads. In the mass chromatogram, the vertical axis indicates intensity (count/sec) and the horizontal axis indicates RT (min). In the mass spectrum, the vertical axis indicates relative intensity (%) and the horizontal axis indicates mass (*m/z*). Compounds are; 14-hydroxy-9-oxo-1,2,3,4,5,6,10,19-octanor-13,17-secoandrostane-7,17-dioic acid (**XVIII**), 9,14-dihydroxy-1,2,3,4,5,6,10,19-octanor-13,17-secoandrostane-7,17-dioic acid (**XXXI**), and 3-hydroxy-6-methyl-7-oxo-decane-1,10-dioic acid (**XXVI**).

